# Data-driven extraction of human kinase-substrate relationships from omics datasets

**DOI:** 10.1101/2022.01.15.476449

**Authors:** Benjamin Dominik Maier, Borgthor Petursson, Alessandro Lussana, Evangelia Petsalaki

**Affiliations:** European Molecular Biology Laboratory, European Bioinformatics Institute (EMBL-EBI), Wellcome Genome Campus, Hinxton, Cambridgeshire, CB10 1SD, United Kingdom

## Abstract

Phosphorylation forms an important part of the signalling system that cells use for decision making and regulation of processes such as cell division and differentiation. To date, a large portion of identified phosphosites are not known to be targeted by any kinase. At the same time around 30% of kinases have no known target. This knowledge gap stresses the need to make large scale, data-driven computational predictions.

In this study, we have created a machine learning-based model to derive a probabilistic kinase-substrate network from omics datasets. Our methodology displays improved performance compared to other state-of-the-art kinase-substrate prediction methods, and provides predictions for more kinases. Importantly, it better captures new experimentally-identified kinase-substrate relationships. It can therefore allow the improved prioritisation of kinase-substrate pairs for illuminating the dark human cell signalling space.

Our model is integrated into a web server, SELPHI_2.0_, to allow unbiased analysis of phosphoproteomics data, facilitating the design of downstream experiments to uncover mechanisms of signal transduction across conditions and cellular contexts.

## Introduction

Cells relay information through signalling networks(1) that are typically regulated by post translational modifications (PTM), with phosphorylation being the best studied one among these(2). Phosphorylation of proteins is catalysed by kinases that target a specific set of substrates, changing their state and/or function. The importance of understanding kinase regulatory networks is highlighted by the fact that a large portion of targeted therapies that are currently used and being developed target kinases(3). Despite this, to date only a small portion (∼5%) of the more than 100,000 known phosphosites have a known upstream kinase(4). At the same time 150 kinases have no known substrate and 90% of the phosphosites are assigned to the 20% of the best studied kinases. Given that several studies have demonstrated that the understudied kinases can be just as important for health as the well-studied ones(4), this points to a bias in the literature, where researchers prioritise well-studied proteins. In addition, publicly available databases, such as KEGG(5), Reactome(6), Omnipath(7) and others, that describe our current knowledge of cell signalling pathways, present a static view that represents the ‘average’ cell and doesn’t capture condition-specific signalling networks. As a result these literature-defined static signalling pathways have a limited explanatory value when it comes to the analysis of phosphoproteomics data.

Mass spectrometry-based phosphoproteomics data sets present relatively unbiased views of a cell’s signalling state and could be used for extracting unbiased signalling networks from specific experimental conditions(8, 9). However, most well-performing methods that have been developed to this end, base their predictions on prior networks((8, 9) in the form of known pathways mentioned above, thus perpetuating existing literature biases. For example, PropheticGranger, one of the top scoring methods in the HPN-DREAM challenge((8, 9), applies heat diffusion coupled with L1 penalized regression(10) on a network based on the Pathway Commons database(11). There is thus a need for a more unbiased prior network to be used in methods for network inference of signalling networks from phosphoproteomics data.

Several methods have been developed to predict kinase-substrate interactions(12). Most of them use in some way either kinase specificity maps in the form of Position Specific Scoring Matrices (PSSMs)(13–16), protein interaction or other types of networks(17), structural information(18) or combinations of these(19). More recently co-expression at the protein level(20) or co-phosphorylation has been used(21). A wide range of approaches is used from basic PSSM scoring, log odds ratios(14), profile Hidden Markov Models(16), to machine learning (neural networks(19), SVMs(22), Bayesian decision theory(23)) and more recently deep or transfer learning(24, 25). However, most of these methods are only able to make predictions for a small fraction of the kinome and many of them only make predictions at the kinase family level.

There have recently become available Position Specific Scoring Matrices (PSSMs) describing the specificity preferences of almost all human kinases (26, 27). This information is invaluable for mapping kinase-substrate relationships but it is difficult to deconvolve kinase-substrate relationships when they have similar PSSMs. Integrating co-expression (at the gene and protein level) and other types of information can help resolve this issue, allowing the generation of kinase-substrate regulatory networks in a data-driven way.

In previous work, we used a machine learning approach, combining predictors such as co-phosphorylation, co-expression and kinase specificity models to derive such a data-driven kinase-kinase regulatory network(28). The resulting network provided predictions only at the protein level, whereas it is known that proteins have multiple functional phosphosites, often with entirely different or even opposing functions. E.g. phosphorylation of Y530 found on the Src kinase leads to its inhibition while dephosphorylation of Y530 and phosphorylation of Y419 leads to its activation(29). In addition, while kinase regulatory networks form the ‘skeleton’ of the phospho-based signalling networks, non kinase substrates are also critical e.g. in the case of adaptor proteins forming scaffolds to regulate signal propagation(30), for phosphatases removing phosphosites to shut signals off, regulation of transcription factor translocation or activities(31) and others.

To address these points, here, we have extended our previous machine learning model introducing additional predictors to include non-kinase substrates and to provide a list of highly probable novel experimentally predicted kinase-substrate relationships(27, 32, 33) at the phosphosite level. Our method, called SELPHI_2.0_ (Systematic Extraction of Linked PHospho-Interactions 2.0), is able to make predictions for less studied kinases and substrates compared to those found in the literature and perform better than established kinase-substrate prediction methods(19, 20, 34–36). To facilitate the use of this network rather than literature-based pathways as a prior network, we created a web server that extracts context-specific signalling networks from global phosphoproteomics datasets in addition to performing other functional analyses. It substantially improves and extends the functionalities of the previously published SELPHI web server(8) and provides an accurate backbone for network extraction rather than a simple correlation measure for kinase-substrate associations that was used by SELPHI. We have made this service available through the url: https://selphi2.com.

## Experimental Procedures

### Generation of kinase-substrate probabilistic network

Information on known kinase-substrate relationships as well as a list of phosphosites and the amino acids sequences surrounding them was gathered from PhosphoSitePlus(37) (Downloaded on 03.09.2024). Functional scores and predictive features on phosphosites were downloaded from Ochoa and colleagues(38) (**Supplemental table 1**). Proteomics data sets for co-regulation were downloaded from Mertins and colleagues(39) as well as Hijazi and colleagues(33). Expression data was gathered from GTEx(40) (Downloaded on 03.09.2024) and the Human Protein Atlas version 23.0(41) (Downloaded on 03.09.2024). Experimentally supported kinase-substrate relationships were downloaded from two recent publications(32, 33).

Predictions were made between 421 kinases, and 238,374 phosphosites found on 17,469 proteins that were listed in PhosphositesPlus(37) and had a functionality score(38) assigned to them.

As a positive training set we used 14,542 kinase-phosphosite relationships extracted from PhosphositePlus(37) (Downloaded on 03.09.2024). As there are no databases with information on known negative kinase-substrate pairs and given the fact that biological networks tend to be sparse, random samples of kinase-substrate relationships ten, twenty, and fifty times as large as the positive set were used as a negative training set. There were no differences between the results, we therefore present those where we used a negative set fifty times as large as the positive set.

We created or acquired 48 features (see below and **Supplemental Table 1**) considering different metrics that could affect a kinase-substrate relationship, from co-regulation in high throughput datasets to match of a phosphosite to a kinase specificity model as described by position-specific scoring matrices (PSSMs). To select the most useful predictors we evaluated the performance of different feature combinations using 100 training/testing datasets. To select the most useful features for kinase-substrate prediction we generated 100 different training sets using our positive set and a set of random kinase-substrate relationships ten times larger than the positive set as a negative set. Random forest(42) classifiers as implemented in *scikit-learn*(*43*) were used to make predictions. The random forest function has various tunable parameters. *Bootstrap* which enables bootstrap samples to be used when building the trees was set to TRUE. The following parameters were tuned by grid search: *max_depth* which sets the maximum depth of the trees, *min_samples_split* which sets the minimum number of samples needed to split an internal node and *n_estimators* which set the number of trees included in the forest. The following parameters were considered: max_depth: [10, 20, 50, 70, 80, 100], *min_samples_split*:[ 8, 10, 12], *n_estimators*: [150, 500, 1000, 1500]. All combinations were tested with the *GridSearchCV()* function as implemented in scikit-learn. We then performed feature ranking with recursive feature elimination with the cross-validation algorithm, as implemented in *scikit-learn*(*43*). We used ten-fold cross-validation to select the best performing feature composition using the area under the ROC curve to evaluate each model. We selected the 45 features to keep in the final model based on majority voting, *i.e.* selecting features included in >50% of the best performing models (**Supplemental Table 1**).

For training the final model we used the random forest algorithm(42) as implemented in the *scikit-learn* python library(43). The model was trained with the positive set and a random sample one hundred times. Parameters were tuned in each run using grid parameter search as described above. Each model was validated by using ten fold cross-validation. In order to balance the training and test set, a stratified K-fold split as implemented in the *scikit-learn*(*43*) python library was used to keep the proportion between negative and positive labels in each split the same. The average probability across the different models’ outputs was then calculated to assign probabilities to kinase substrates.

To further evaluate the performance of our model on novel kinase-substrate relationships we used predictions from two recently published experimental studies(32, 33) and tested whether these relationships were assigned a higher probability by our method. To quantify the predictive power of our model in each case, the area under the ROC curve was calculated as implemented in the *ROCR* package(44).

### Comparison with other methods

To compare SELPHI_2.0_ with other state-of-the-art peer reviewed methods we assessed how well our method predicted known annotated kinase-substrate relationships (see positive/negative training sets described above) and also novel kinase-substrate relationships supported by experimental procedures. These were derived from the recent works of Sugiyama and colleagues, and Hijazi and colleagues (32, 33), excluding kinase-substrate phosphorylation relationships found in PhosphoSitePlus, to limit literature-based biases.

We compared our method with five other established and most commonly used methods: PhosphoPICK(20), GPS v.5.0(36), GPS v.6.0(24), KinomeXplorer(19), NetPhos v.3.1(34) and LinkPhinder(35). PhosphoPICK and NetPhos v3.1 are available as web servers and sequences of substrate proteins were uploaded onto the servers. For both PhosphoPICK and NetPhos v3.1 we selected no significance threshold to include all predictions. For NetPhos v3.1 we selected all available residues: Serine, threonine and tyrosine. GPS was downloaded from (http://gps.biocuckoo.cn/download.php) and batch kinase prediction was run on Ubuntu 22.04. Confidence threshold was set to “All” to include all predictions. The KinomeXplorer prediction software was downloaded and predictions were made for the phosphosites included in our network. All 11,581,940 LinkPhinder predictions were downloaded from (https://linkphinder.insight-centre.org/download).

Methods were compared by assessing their ability to capture known kinase substrate relationships from PhosphoSitePlus. As the published methods were able to make predictions for different subsets of kinases, we restricted the comparisons to those predictions that were possible by both methods(**Supplemental Table 2**). For each method under comparison, we generated 100 training sets made from the positive set and a negative set selected from random kinase substrate relationships that were present both in SELPHI_2.0_ as well as the method that was under comparison. The negative set was ten times larger than the positive set.

The ability of SELPHI_2.0_ to discriminate between known interactions and interactions with unknown status was then compared to each of the other methods by calculating the average AUC achieved by SELPHI_2.0_ and the method under comparison for the 100 different training sets only including kinase substrate relationships shared by both methods.

In order to estimate the methods’ ability to capture novel kinase substrate interactions we used a set of experimental kinase substrate predictions made previously by Sugiyama and colleagues^8^. For each method, the overlapping predictions between SELPHI_2.0_ and the method under comparison were assessed by how well they captured the experimentally predicted kinase substrate relationships as measured by the AUC. AUC scores were calculated using the *ROCR*(*44*) package.

### Evaluation of model fit to phosphoproteomic data

To capture context specific signalling networks, we constructed a reference SELPHI_2.0_ signalling network by linking the kinase-substrate predictions made in this work to a backbone of a probabilistic kinase kinase regulatory network that we had previously published(28). A probability cutoff of 0.5 was applied on both networks to select high confidence edges.

High-throughput mass spectrometry-based phosphoproteomic data that had been compiled and analysed by Ochoa and colleagues with 436 conditions(45) was fitted to our network by means of the PCSF algorithm(46). We tried all combinations of the of the following PCSF parameters: For the tree parameter, b, we tried fitting anything from 1 to 10 trees to the data, for parameter w or the node tuning we tried the following values: 0.25, 0.5, 0.75, 1.0, 1.25, 1.5 and for the edge tuning, μ, we tried 0.000005, 0.00005, 0.0005, 0.005, 0.05. Edge probabilities subtracted from 1 were used as edge costs. These parameter combinations generated 436 sub networks that gave the highest F1 score. To evaluate the performance of the fitting, the F1 score of kinase-substrate phosphosite relationships retained after the fitting was compared with those that were used in the input.

### Counting number of citations related to proteins

To count the number of citations related to kinases and their substrates we used the Entrez.elink() function from the Biopython module(47). We searched for related articles in the Pubmed database. Linked publications were retrieved from NCBI Entrez Gene(48) database and publications that mention more than ten kinases were filtered out.

### Implementation of SELPHI2.0 web server

The SELPHI_2.0_ web server has been implemented using the *shiny*(*49*) R package. Enrichment analyses on the server are performed using the *enrichR*(*50*) and the clusterProfiler(51) enrichment package and users will be able to select gene sets for enrichment from various databases: Reactome(6), KEGG(5), GO(52) and DISEASES(53). For phosphopeptide clustering, the two clustering algorithms available are Gaussian mixture models as implemented by *mclust*(*54*) and k-means clustering(55). If k-means clustering is selected, silhouette index(56) is used to calculate the optimal number of clusters while *mclust* uses Bayesian information criterion to select the best fitting model(57). The output is presented as a dot plot where the colour indicates the sum of the log_2_ ratios of the phosphosites involved in the enrichment term and size the odd ratio of the enrichment term.

For the calculation of kinase activities the web server uses the 95^th^ percentile of kinase-substrate probabilities to select high confidence kinase-substrates. We then use the Kolmogorov-Smirnov statistical test to assess if the predicted substrates are overrepresented at the upper end of the log_2_ ratio distribution for a given condition. The negative log_10_ transformation of the derived p-value is used to quantify activities. P-values are then multiplied by the average log_2_ ratios of the predicted kinase-substrates. For comparison kinase activities using the PhosphositePlus(37) kinase-substrate data set are also calculated.

## Results

### Generation of a probabilistic human kinase-substrate regulatory network

To create a probabilistic network of kinase-substrate relationships we used the random forest algorithm to combine various predictive variables ranging from co-phosphorylation and co-expression in large datasets, to kinase specificities and features related to the functionality of the phosphosites (**Experimental Procedures**). After running the prediction algorithm 100 times we achieved an average AUC of 0.97 (**Figure 1A**). Of the 73,207,860 predictions made, 721,707 edges were high confidence (probability > 0.5). Of those, around 1.7% were found in PhosphoSitePlus(37) while more than 90% of the known interactions found in our network were assigned a high (> 0.5) probability.

**Figure 1.**
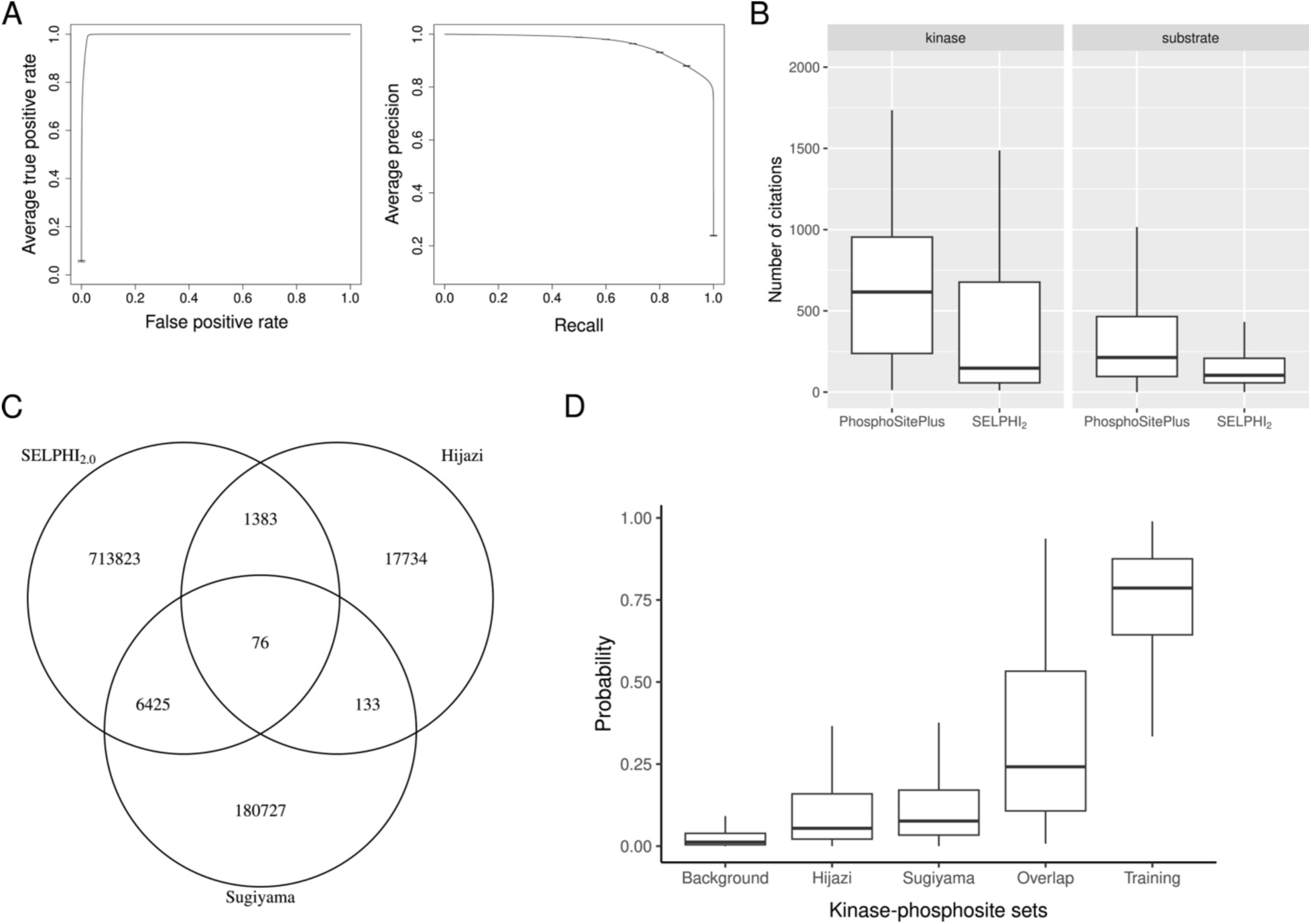
Precision and recall and different cutoffs. (A). The predictive power of our predictive method. AUC under the ROC curve of 0.97 after 100 runs of cross-validation with a random set of negatives and a positive set from phosphosite plus. Precision recall curve for the same set can be seen on the right (B). We found that our classifier is able to make high confidence predictions for less cited kinases and less cited substrates compared to kinase-substrates found in PhosphositePlus (C). We identified 76 high quality kinase-substrate interactions found by SELPHI_2.0_ as well as the two experimental datasets in our study (D). Experimentally validated kinase-substrate relationships were assigned a higher probability compared to the background with kinase-substrate predictions made by both methods being assigned the highest probabilities of all sets.

Importantly, our network provides high confidence predictions covering the ‘dark’ or less well studied human signalling network. Specifically, kinase-substrate relationships found in PhosphositePlus have a median number of 685 citations per phosphorylating kinase and 232 per substrate respectively, whereas our high confidence predictions the median number of citations are 194 for the kinases and 96 for the substrates. We provide proportionally more predictions between kinases and substrates with significantly lower number of citations per protein on average (Kinases: W = 3.37 × 10^9^ , p < 2.2 × 10^-16^ , one-sided Wilcoxon test. Substrate proteins: W= 3.34 × 10^9^ , p < 2.2 × 10^-16^ , one-sided Wilcoxon test)(**Figure 1B**). These predictions include substrates that have not been mentioned in the literature before and have no known upstream kinase, highlighting the value of this network as a reference to explore the less studied part of the phospho-signalling network.

### SELPHI2.0 can be used to identify novel high confidence kinase-substrate relationships

To assess the ability of our method to predict novel kinase-substrates, we looked at how well we discerned between a set of experimental kinase-substrate predictions not hitherto found in the literature and the rest of the unsupported predictions. To this end, we used experimentally predicted kinase-substrate relationships from two recent papers(32, 33). In short, one study introduces kinases to dephosphorylated peptides from cell lysis, while the other predicts kinase-substrates based on changes in phosphorylation following kinase inhibition.

We found that the overlap between the three sets, the two experimental sets and the SELPHI_2.0_ predictions, was relatively low (**Figure 1C**). Due to the complementary characteristics between the two experimental methods we reasoned that kinase-substrate relationships identified by both studies should have higher levels of confidence. Our method assigned significantly higher probabilities to experimentally supported edges compared to the rest of the network (**Figure 1D**). Furthermore, we found that kinase-substrate relationships supported by edges found in both data sets had an even higher probability assigned to them (**Figure 1D**) compared to edges supported by either.

### SELPHI2.0 outperforms the state-of-the-art methods for kinase-substrate prediction

In comparison to 5 state-of-the-art kinase-substrate prediction methods (PhosphoPICK(20), GPS v.5.0(36) & v.6.0(24), KinomeXplorer(19), NetPhos v.3.1(34) and LinkPhinder(35)), SELPHI_2.0_ performs generally better on identifying known kinase-substrate interactions (**Supplemental Figure 1**), while making predictions for more kinases. When comparing the performance of these methods using an independent dataset of experimentally supported kinase-substrate pairs(32, 33), having removed those that already exist in PhosphositePlus to remove the effect of literature bias on the methods, we found that our network performs better than the others with GPSv.6.0 performing the closest with AUC=0.81 compared to our method’s 0.83. NetPhos v3.1 achieved AUC of 0.69 compared to SELPHI_2.0_’s AUC of 0.75, but made predictions for only 17 (38 kinase genes) kinases. KinomeXplorer(19) achieved AUC of 0.61 compared to SELPHI_2.0_’s AUC of 0.81 for the same set (**Supplemental table 2**). Other methods performed close to random (**Figure 2**).

**Figure 2:**
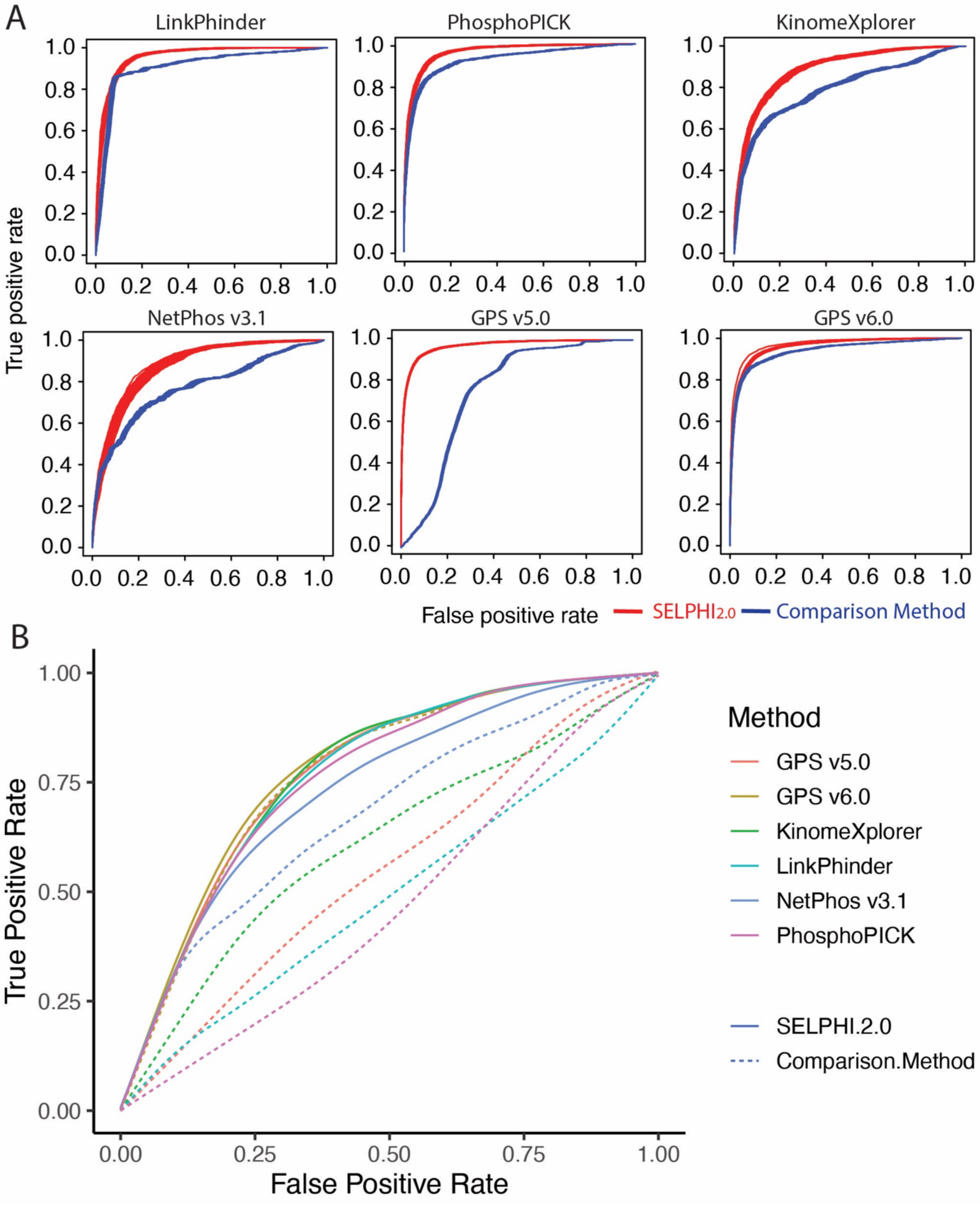
Comparison between SELPHI_2.0_ (solid lines) and other state of the art kinase-substrate prediction methods (dashed). The ability of each method to discern between test (A) or experimentally supported (B) kinase-substrate relationships and the rest of the relationships unsupported by the experiments. In all cases SELPHI_2.0_ performed better than the state of the art methods.

### Extraction of dataset-specific networks from our reference network selects for known interactions

We fitted our network to a set of mass spectrometry-based global phosphoproteomic data sets generated under different conditions. We used a compilation of mass spectrometry data sets compiled and reanalyzed downloaded from an earlier publication(45). To fit our predictions to high-throughput data, the predicted kinase-substrate edges were combined with a kinase-kinase regulatory network that we had generated previously(28), forming a network of kinase-kinase regulatory relationships with phosphosites as nodes without outgoing edges (**Experimental Procedures**). In order to select high confidence edges, we used edge probability of 0.5 as a threshold for both the kinase-kinase regulatory network and the kinase-substrate predictions.

To fit the combined network to the high-throughput data sets, we used Prize collecting Steiner’s forest as implemented in the R package PCSF(46). We found that by optimising the edge cost against the node prizes we were able to select for edges found in the literature. The F1 scores of the fitted subnetworks (n = 415) were 0.19 compared to the unpruned input with a F1 score of 0.031. The improvement in precision was even greater with the mean precision of the pruned subnetworks being 0.22 while the precision was 0.016 for the unpruned input networks. Both comparison yielded a significant difference (F1 score: W= 2.14 × 10^6^, p < 2.2 × 10^-16^ , precision: W = 2.14 × 10^6^, p < 2.2 × 10^-16^), indicating this combination of kinase-kinase regulatory network and kinase-substrate predictions can be used to extract high probability context-specific subnetworks (**Figure 3**).

**Figure 3:**
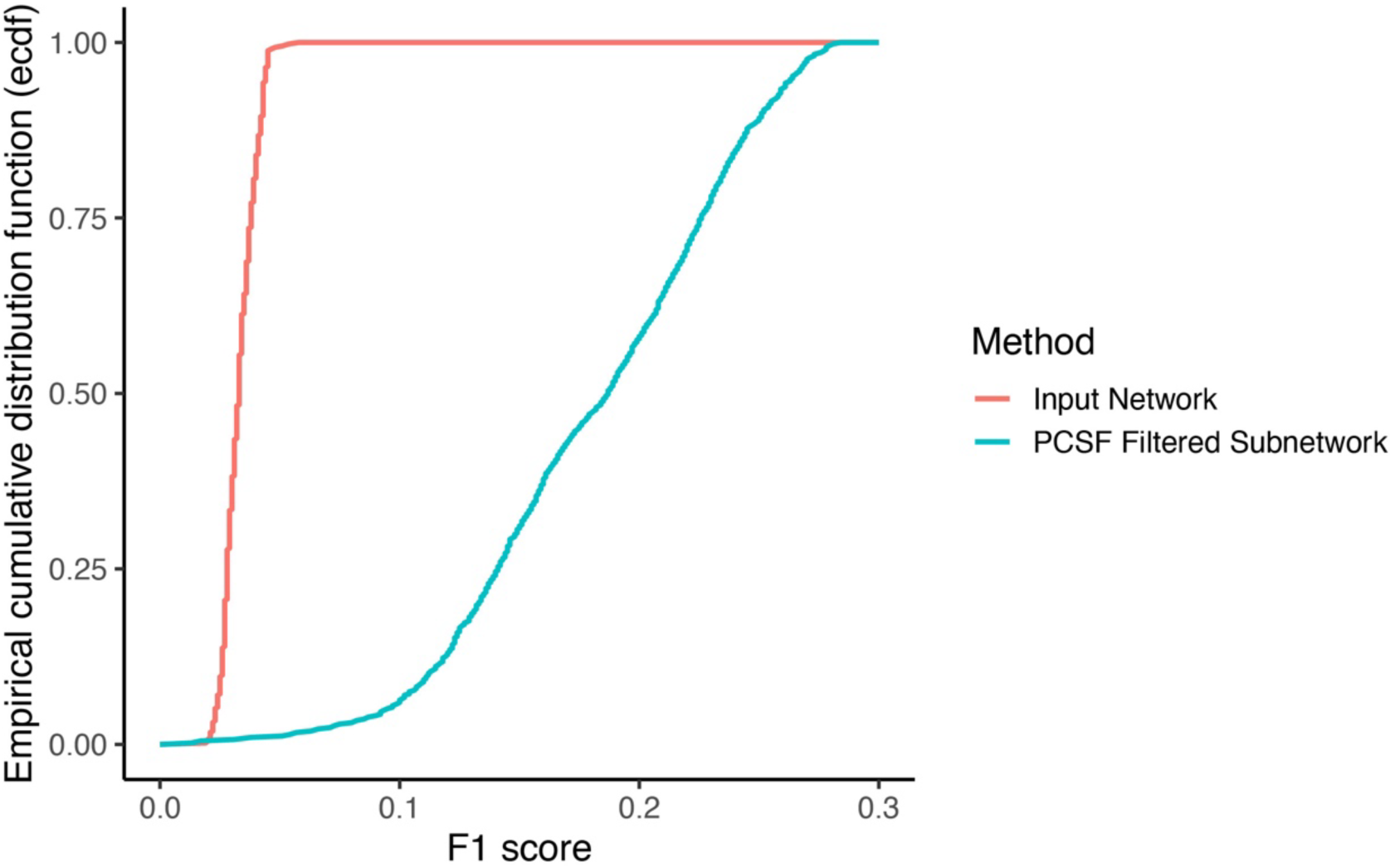
Fitting the kinase-substrate predictions to experimental data selects for known kinase-substrate relationships with median F1 score of 0.031 for the unfitted sub network compared to 0.19 for the fitted input.

### SELPHI_2.0_ web server

We have created a user-friendly web server, SELPHI_2.0_, that allows less biased analyses of global phosphoproteomics datasets, to help users shed light on the mechanisms underpinning their data, including context-specific and understudied signalling processes. The basis of this web server is the data-driven network of kinase substrate predictions described above. SELPHI_2.0_ improves on an earlier resource SELPHI (Systematic Extraction of Linked Phospho-Interactions)(8), which calculated kinase-substrate associations based on correlations and filtering based on prior knowledge.

Users can use SELPHI_2.0_ to retrieve the top kinase-substrate predictions for the phosphosite included in their data and extract condition specific kinase activity profiles (**Figure 4A**). They can also perform standard functional enrichment analysis either on their up/down-regulated phosphosites or on clusters of phosphosites identified through the server. Furthermore, users can identify the active signalling sub-networks in their data by mapping it to our SELPHI_2.0_ reference kinase-substrate network (**Experimental Procedures; Figure 4B**). Finally, the server also provides higher confidence relationships by overlapping our predictions with the two experimentally-derived data sets published previously(32, 33) . This way the user can prioritise edges that have been corroborated by independent experimental results as well as large scale omics datasets (**Figure 4C**).

**Figure 4:**
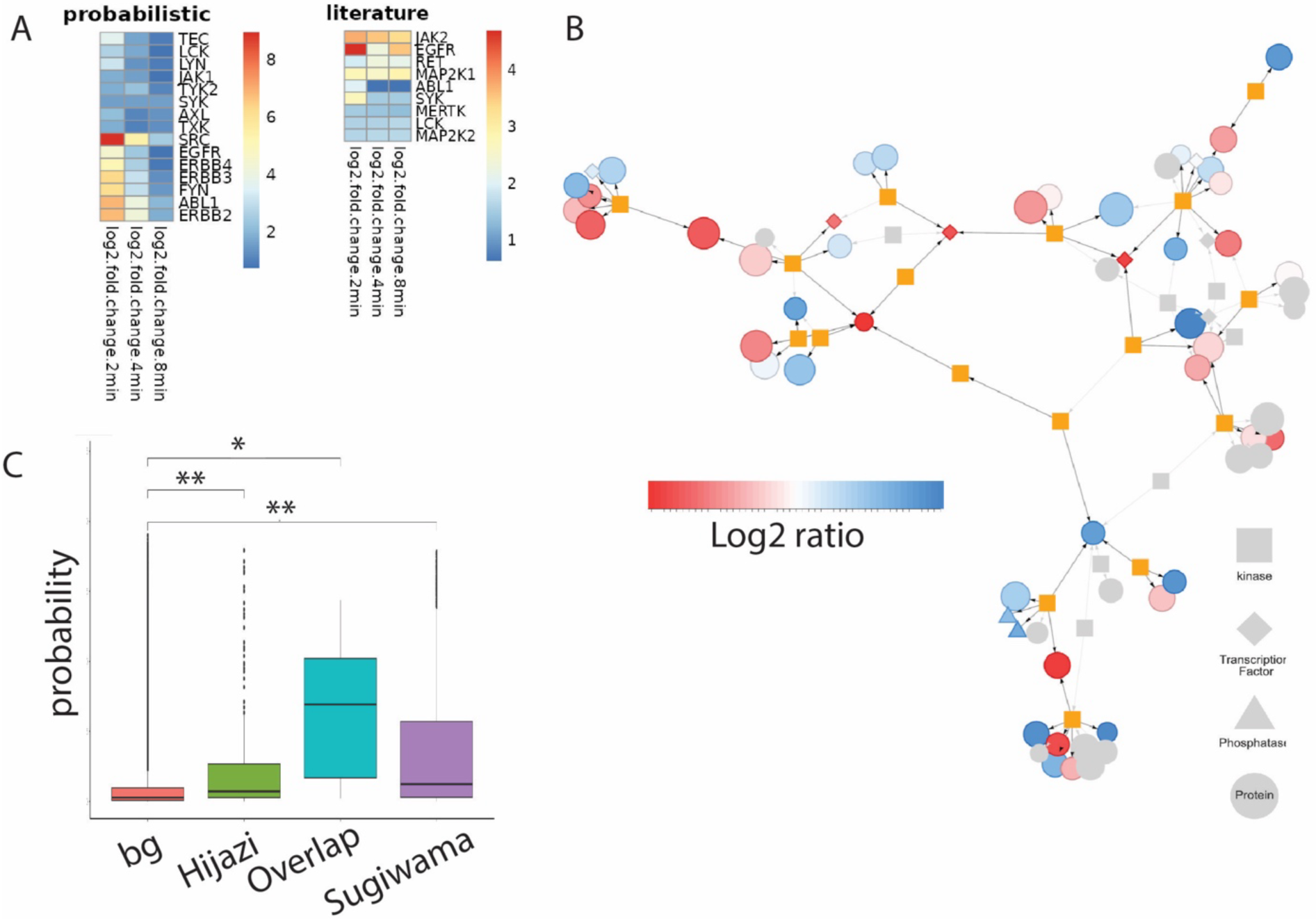
Examples of the SELPHI_2.0_ web server outputs using as input the first example in the web server. The user can upload phosphoproteomics data and fit the SELPHI_2.0_ reference network to it (A). Kinase activities across the different conditions or time points can be estimated based on kinase substrate predictions (Heatmap titled Prob_Y) and kinase substrates as listed in PhosphoSitePlus (Heatmap, titled PPS_Y) for comparison (B). The user can assess if experimentally predicted edges from independent studies corroborate the prediction made by SELPHI_2.0_. Furthermore, The user can download a list of these experimentally corroborated edges (C).

## Discussion

Kinase-substrate networks form the backbone of cell signalling responses and are critical for cell function in health and disease(58). It is, therefore, important to accurately annotate kinase-substrate relationships, but the vastness of the potential kinase-substrate interaction space makes computational means necessary for their prioritisation. Here, we have used a machine learning model, integrating information largely from high throughput datasets, to generate a probabilistic human kinase-substrate network at the phosphosite level that includes 421 kinases and 238,374 phosphosites.

Our method, called SELPHI_2.0_, performs better than the five state-of-the-art methods tested(19, 20, 34–36), not only on the benchmark set but importantly on entirely new experimentally supported data(32, 33). This is true despite including predictions for more kinases than all other methods. This suggests that our network is a good starting point for prioritisation of new kinase-substrate relationships. NetPhos(34) proved to be an exception, performing well both at discerning between known positives and negatives as well as experimentally validated edges. It should be kept in mind though that NetPhos only makes predictions for 17 kinases (38 kinase genes). When overlaying our predictions with the two experimentally supported datasets, the resulting 24 high confidence kinase-substrate relationships are supported by both *in vitro* and cell line-based experimental data, and our data-driven machine learning approach, giving them particularly high confidence.

Our predictors, with the exception of the literature-derived PSSMs, were based on unbiased, high throughput datasets including global phosphoproteomics datasets. This, in combination with the coverage of the dark space that our method affords, is a step towards reducing the bias in cell signalling studies. Our network provides much wider coverage of the kinase signalling space than current knowledge and most other available kinase-substrate prediction methods and is more accurate.

With the median citation number for kinases and substrates in the network being 3.5 and 2.4 times less than the PhosphositePlus database, our network forms a springboard for the exploration of the dark human cell signalling space. Given the importance of a prior network in methods of signalling network inference, this network will significantly contribute to a better understanding of the role of the understudied space in the context of our current knowledge, and will allow methods to generate networks that more accurately reflect the data.

Our method combines the power of PSSMs for kinase-substrate association mapping based on kinase specificity information(26, 27), with functional information from unbiased functional genomics datasets. This can open the door to improved kinase activity estimation from phosphoproteomics datasets improving our ability to describe context-specific signalling processes. For example, tools like PhosX(59)use kinase specificity scores rather than known substrates, can readily take advantage of the SELPHI_2.0_ scores to calculate kinase activities considering both the PSSMs and multiple additional metrics describing the kinase substrate relationships, such as coexpression and others.

We have created a web server called SELPHI_2.0_, which allows the easy analysis of global phosphoproteomics data to design downstream experiments. Global phosphoproteomics datasets provide a unique snapshot of the signalling networks that are active in a given cell type and condition, i.e. context. It is, however, very difficult to extract unbiased ‘active’ signalling networks from these datasets largely due to the inherent noisiness of both the experimental measurements and cell signalling itself, and due to the sparsity of these datasets(4). In addition to standard analyses such as functional enrichment and clustering, it provides a method for extracting context specific networks from the data. SELPHI_2.0_ improves on a previous web server, SELPHI, which was using simple correlation between kinases and potential substrates filtered by prior knowledge information for the generation of context-specific signalling networks from phosphoproteomics data. It thus perpetuated the literature bias, for kinases with existing knowledge, while it provided very noisy associations for the understudied ones.

In SELPHI_2.0_ the user can derive dataset-specific networks extracted from our reference probabilistic network using PCSF. As signalling responses vary depending on the conditions and cell properties (e.g. tissue, state etc) and current signalling pathways poorly explain observed phosphoproteomics datasets, methods that infer context-specific signalling networks are necessary to understand signalling in specific contexts of interest. In the future, more sophisticated statistical methods, such as MRA(60) or Integer Linear Programming methods(61), can be implemented as additional options.

Accumulation of false positives is a persistent problem when kinase-substrates are predicted. This is partly due to the large fraction of the signalling network that is currently unknown or understudied but a fraction of these predictions can be assumed to be false positives, even though this can’t possibly be confirmed. This is likely true for our method as well and users of the network should take this into consideration when interpreting the results of their analysis. Nevertheless, the vast understudied signalling space requires computational approaches for prioritisation of hypotheses and the performance of our network compared to the state of the art indicates that it provides a good starting point. As more high quality high throughput phosphoproteomic data sets become available our model can be further improved.

We expect that SELPHI_2.0_ will allow the improved prioritisation of kinase-substrate pairs for illuminating the dark human cell signalling space, both through smaller scale signalling studies and through acting as a relatively unbiased prior in signalling network inference approaches. Our web server also makes this network accessible to non bioinformatics experts and allows the interrogation of large scale phosphoproteomics datasets in an unbiased and context-specific way. We thus expect it to facilitate the design of downstream experiments to illuminate mechanisms of signal transduction across conditions and contexts.

## Data and code availability

The SELPHI_2.0_ web server is freely available under: https://selphi2.com and all code for reproducing this project can be found at https://gitlab.ebi.ac.uk/petsalakilab/selphi_2.0 and https://github.com/alussana/selphi2.0-pssms. A Docker image of the web server can be found at: https://gitlab.ebi.ac.uk/petsalakilab/selphi_2. The feature table, positive training set. Prediction results and citation matrix can be found in Zenodo: https://zenodo.org/records/13821321?preview=1&token=eyJhbGciOiJIUzUxMiJ9.eyJpZCI6IjIyYjliMThhLWJjNTQtNDMwYy05NTJkLWNiM2E1NjA0MDFkYiIsImRhdGEiOnt9LCJyYW5kb20iOiJiZDk3NmU1MjY4MWEwY2RlOTg4MGJmMmM3YWFkODNhMSJ9.T_-xpVVN5MVVHVTM5xH6YXXcfTZHI94gFIJb92DIAxwtOzCEx5YZXVP_I8o79c9XI72qV9-MxDtsbdMD1t5thg

## Author Contributions

BP: Conceptualization, Methodology, Validation, Formal analysis, Investigation, Data curation, Writing - Original Draft, Visualization. BM & AL: Methodology, Validation, Formal analysis, Investigation, Software, Data curation, Writing - Review & Editing. EP: Conceptualization, Methodology, Resources, Writing - Original Draft, Writing - Review & Editing, Supervision, Project administration, Funding acquisition.

## Acknowledgements

This work was supported by the European Molecular Biology Laboratory (EMBL), and EMBL-EBI. BP, BM and AL were funded by the EMBL International PhD Programme. EMBL-EBI IT Support is acknowledged for provision of computer and data storage servers.

