## Supplementary Tables for "Data-driven extraction of human kinase-substrate relationships from omics datasets"

**Running Title:** Prediction of human kinase-substrate relationships

### Supplementary information table of contents

Supplemental Table 1

Supplemental Table 2

**Supplemental Table 1. Features used in the development of the machine learning model to predict human kinase-substrate relationships.**

| Name of feature | Description | Imputation |
|---|---|---|
| Residue | Indicator of phospho acceptor residue: S/T/Y | No missing values |
| PWM score | Score generated by fitting phosphosites to PWMs described in previous publication(28). | 0 |
| Coreg_293 | Spearman's correlation between estimated kinase activities and target phosphosite's phosphorylation(39). Kinase activities were estimated with the KSEA algorithm(45) | 0 |
| Functional_score | Score indicating the probability of phosphosite to be functional(38, 45). | No missing values |
| GTEx | Co-expression between kinase and putative substrate genes across tissues(40). Co-expression was calculated using Spearman's correlation. | 0 |
| RNA_tissue | Co-expression between kinase and putative substrate gene across tissues(41). Co-expression was calculated using Spearman's correlation | 0 |
| RNA_cell | Co-expression between kinase and putative substrate gene across cell lines(62). Co-expression was calculated using Spearman's correlation | 0 |
| Kinase_selectivity | Skewness of kinase expression distribution across different tissues(41). Skewness was calculated by using the e1071 R library to calculate the skewness (https://CRAN.R-project.org/package=e1071). | 0 |

| | | |
|---|---|---|
| Substrate_selectivity | Skewness of substrate gene expression distribution across different tissues(41). Skewness was calculated by using the e1071 R library to calculate the skewness (https://CRAN.R-project.org/package=e1071). | 0 |
| NTERA2_coreg | Spearman's correlation between estimated kinase activities and target phosphosite's phosphorylation across 63 inhibitor conditions in NTERA2 cell line(33). Kinase activities were estimated with the KSEA algorithm(45). | 0 |
| MCF7_coreg | Spearman's Correlation between estimated kinase activities and target phosphosite's phosphorylation across 63 inhibitor conditions in MCF7 cell line(33). Kinase activities were estimated with the KSEA algorithm(45). | 0 |
| HL60_coreg | Spearman's correlation between estimated kinase activities and target phosphosite's phosphorylation across 63 inhibitor conditions in HL60 cell line(33). Kinase activities were estimated with the KSEA algorithm(45). | 0 |
| Is_DISOPRED | Indicates if phosphosite is disordered (DISOPRED score(63) > 0.5). This feature was downloaded from previous publication by Ochoa and colleagues(38). | Phosphosites with missing values are assumed to be disordered |
| DISOPRED score | DISOPRED score of target phosphosite(63). This feature was downloaded from previous publication by Ochoa and colleagues(38). | Median imputation |
| Exp3d_ala_ddG_effect | Discretized changes in Gibb's energy upon mutation from original residue to alanine. Measured by FoldX v4(64). This feature was downloaded from previous publication by Ochoa and colleagues(38). | Missing data set to unknown |

| | | |
|---|---|---|
| Exp3d_acid_ddG_effect | Discretized average changes in Gibb’s energy upon mutation from original residue to acidic residue Measured by FoldX v4(64).. This feature was downloaded from previous publication by Ochoa and colleagues(38). | Missing data set to unknown |
| Log_10_of_hotspot_pval_min | Quantification of the prevalence of phosphorylation events within the structural region. The p-value indicates the enrichment of phosphosites within the region in question(65). This feature was downloaded from previous publication by Ochoa and colleagues(38). | 0 |
| Is_hotspot | Indicates If the phosphosite has assigned hotspot enrichment value of <3.36e-07 and is present in more than 10 MS data sets (65). This feature was downloaded from previous publication by Ochoa and colleagues(38). | Missing values set to FALSE |
| Is_Interface | Experimentally resolved or modelled interaction interfaces were obtained from Interactome3d(66). NACCESS(67) was used to calculate relative solvent accessibility of atoms. In cases where the relative solvent accessibility differed between interacting and non-interacting form, the residue was considered to be found on the interface. This feature was downloaded from a previous publication by Ochoa and colleagues(38). | Missing values are set to FALSE |
| Adj_ptms w21 | Number of phosphosites within +/- 10 residue window on either side of the target phosphosite. This feature was downloaded from a previous publication by Ochoa and colleagues(38). | Zero |
| Netpho_max_all | Highest posterior probability obtained from all NetPHOREST v2.1(19) models. This feature was downloaded from a previous publication by Ochoa and colleagues(38). | Median imputation |

| | | |
|---|---|---|
| Netpho_max_KIN | Highest posterior probability obtained from all kinase models found in NetPHOREST v2.1(19). This feature was downloaded from a previous publication by Ochoa and colleagues(38). | Median imputation |
| Paxdb_abundance_log10 | Consensus protein abundance downloaded from the PaxDb(68) database. This feature was downloaded from a previous publication by Ochoa and colleagues(38). | Median imputation |
| W0_mya | Age of the target phosphosite. This feature was downloaded from a previous publication by Ochoa and colleagues(38). | 0 |
| W3_mya | Age of the +/- 3 residue region surrounding the target phosphosite. This feature was downloaded from a previous publication by Ochoa and colleagues(38). | 0 |
| Quant_top1 | Number of times phosphosite was found within the top 1% of regulated sites across 435 conditions(45). This feature was downloaded from a previous publication by Ochoa and colleagues(38). | 0 |
| Quant_top5 | Number of times phosphosite was found within the top 5% of regulated sites across 435 conditions(45). This feature was downloaded from a previous publication by Ochoa and colleagues(38). | 0 |
| PWM_max_mss | The best fit to one of the PSSMs of 143 different kinases known to have at least 10 known substrates as measured by the MATCH algorithm(69). This feature was downloaded from a previous publication by Ochoa and colleagues(38). | Median imputation |
| ACCpro | Prediction of solvent accessibility with ACCpro(70). This feature was downloaded from a previous publication by Ochoa and colleagues(38). | Missing values set to unknown |
| SSpro | Prediction of secondary structure with SSpro(70). This feature was downloaded from a previous publication by Ochoa and colleagues(38). | Missing values set to unknown |
| SSpro8 | Indicates the class of secondary structure as predicted by SSpro8(70). This feature was downloaded from a previous publication by Ochoa and colleagues(38). | Missing values set to unknown |

| | | |
|---|---|---|
| SIFT_min_score | Conservation using SIFT[26] score as a proxy. This feature contains the minimum score across all variants. This feature was downloaded from a previous publication by Ochoa and colleagues(38). | Median imputation |
| SIFT_mean_score | Conservation using SIFT score as a proxy(71). This feature contains the mean score across all variants. This feature was downloaded from a previous publication by Ochoa and colleagues(38). | Median imputation |
| SIFT _ala_score | Conservation using SIFT(71) score as a proxy. This feature contains the score of alanine variants. This feature was downloaded from a previous publication by Ochoa and colleagues(38). | Median imputation |
| SIFT_acid_score | Conservation using SIFT(71) score as a proxy. This feature contains the average score of variants leading to negative charge. This feature was downloaded from a previous publication by Ochoa and colleagues(38). | Median imputation |
| IsProteinDomain | This feature indicates if the target phosphosite is found within a protein domain(72). This feature was downloaded from a previous publication by Ochoa and colleagues(38). | Missing values set to FALSE |
| IsProteinKinaseDomain | This feature indicates if the target phosphosite is found within a kinase domain(72). This feature was downloaded from a previous publication by Ochoa and colleagues(38). | Missing values set to FALSE |
| IsUniprotRegion | This feature indicates if the target phosphosite is found within any other curated motif(72). This feature was downloaded from a previous publication by Ochoa and colleagues(38). | Missing values set to FALSE |
| IsCytoplasmic | This feature indicates if the target phosphosite is found within a cytosolic region of a transmembrane protein(72). This feature was downloaded from a previous publication by Ochoa and colleagues(38). | Missing values set to FALSE |
| IsMotif | This feature indicates whether the target phosphosite is found within any other curated motif(72). This feature was downloaded from a previous publication by Ochoa and colleagues(38). | Missing values set to FALSE |

| | | |
|---|---|---|
| IsELMLinearMotif | This feature indicates if the flanking region of the phosphosite is listed under linear motifs (CLV, DEG, DOC, MOD, TRG, and/or kinase motif)) in ELM(73). This feature was downloaded from a previous publication by Ochoa and colleagues(38). | Missing values set to FALSE |
| IsEV_ala_prediction_epistatic5 | This feature indicates if alanine mutation leads to epistatic effects as calculated with the EVmutation algorithm(74). This feature was downloaded from a previous publication by Ochoa and colleagues(38). | Missing values set to FALSE |
| IsKinaseCoreg | This feature indicates if the target phosphosite is coregulated with a kinase. Derived from kinase activities estimated in previous publication[12]. This feature was downloaded from a previous publication by Ochoa and colleagues(38). | Missing values set to FALSE |
| serThrPssmScore | The transformed PSSM score between the given kinase-phosphosite pair. The raw PSSM score(27) is transformed between 0 and 1 based on the quantile it falls in, considering a kinase-specific background distribution of proteome-wide raw PSSM scores. These background scores are computed on a set of more than 200,000 human phosphosites annotated in the PhosphositePlus database(37) (September 2023). Phosphosites with raw PSSM score equal to 0 are discarded, and the remaining are used to determine the values of the 10,000-quantiles of the raw PSSM score distribution. | 0 |
| tyrPssmScore | This feature is computed exactly as serThrPssmScore, but using the PSSMs sourced from(26) | 0 |
| | Excluded Features | |
| *IsUniprotRepeat* | *This feature indicates if the target phosphosite is found within a repeated motif(72). This feature was downloaded from a previous publication by Ochoa and colleagues(38).* | *Missing values set to FALSE* |

| | | |
|---|---|---|
| *IsUniprotZnFinger* | *This feature indicates if the target phosphosite is found within a zinc finger(72). This feature was downloaded from a previous publication by Ochoa and colleagues(38).* | *Missing values set to FALSE* |
| *IsUniprotCompBias* | *This feature indicates if the target phosphosite is found within a compositionally biased region(72). This feature was downloaded from a previous publication by Ochoa and colleagues(38).* | *Missing values set to FALSE* |

**Supplemental table 2. Overview over the number of kinase substrate relationships, kinases and the size of the positive set as well as the overlap with Sugiyama et al.**

| **Method** | **Number of supported kinases** | **Number of kin-substrates shared with SELPHI2.0** | **Number of kin-substrates shared with SELPHI2.0 and predicted by Sugiyama et al.** |
|---|---|---|---|
| PhosphoPICK | 107 | 7,006,604 | 17,060 |
| GPS v5.0 | 479 | 18,319,104 | 65,141 |
| GPS v6.0 | 349 | 25,173,556 | 71,882 |
| NetPhos v3.1 | 17 | 5,528,807 | 13,116 |
| LinkPhinder | 327 | 1,343,230 | 4,568 |
| KinomeXplorer | 193 | 7,750,644 | 30,645 |
